# The Parkinson’s Disease GWAS Locus Browser

**DOI:** 10.1101/2020.04.01.020404

**Authors:** Francis P. Grenn, Jonggeol J. Kim, Mary B. Makarious, Hirotaka Iwaki, Anastasia Illarionova, Kajsa Brolin, Jillian H. Kluss, Artur F. Schumacher-Schuh, Hampton Leonard, Faraz Faghri, Kimberley Billingsley, Lynne Krohn, Ashley Hall, Monica Diez-Fairen, Maria Teresa Periñán, Cynthia Sandor, Caleb Webber, J. Raphael Gibbs, Mike A. Nalls, Andrew B. Singleton, Sara Bandres-Ciga, Xylena Reed, Cornelis Blauwendraat, on behalf of the International Parkinson’s Disease Genomics Consortium (IPDGC)

## Abstract

Parkinson’s disease (PD) is a neurodegenerative disease with an often complex genetic component identifiable by genome-wide association studies (GWAS). The most recent large scale PD GWASes have identified more than 90 independent risk variants for PD risk and progression across 80 loci. One major challenge in current genomics is identifying the causal gene(s) and variant(s) from each GWAS locus. Here we present a GWAS locus browser application that combines data from multiple databases to aid in the prioritization of genes associated with PD GWAS loci. We included 92 independent genome-wide significant signals from multiple recent PD GWAS studies including the PD risk GWAS, age-at-onset GWAS and progression GWAS. We gathered data for all 2336 genes within 1Mb up and downstream of each variant to allow users to assess which gene(s) are most associated with the variant of interest based on a set of self-ranked criteria. Our aim is that the information contained in this browser (https://pdgenetics.shinyapps.io/GWASBrowser/) will assist the PD research community with the prioritization of genes for follow-up functional studies and as potential therapeutic targets.

## Introduction

Parkinson’s disease is a multifactorial disease where both genetic and environmental risk factors play a role. In the past decade, approximately 20 genes have been associated with PD or parkinsonism in families [1]. Over 90 common variants have been associated with the sporadic PD risk, onset and progression using genome-wide association studies (GWAS) [2–4].

One major challenge remaining after GWAS identification of risk loci is the localization and characterization of specific causal variant(s) and gene(s) at each locus. A common misconception is that the most significant GWAS variant exerts an effect on the nearest gene (as commonly reported in GWAS manuscripts), but this is unlikely to be the case. First, the GWAS variant is not necessarily causative on its own, but is instead likely to tag a functional region or variant in high linkage disequilibrium (LD). Second, variants in noncoding regions containing regulatory sequences may impact distant genes by altering the three dimensional chromatin conformation and therefore those genes may be located outside of the predefined LD region [5,6]. Several different approaches can be taken when prioritizing genes for each significant variant. For PD, these approaches include using single cell RNA-seq to determine gene expression in relevant cell populations [7], transcriptome-wide association studies [8], and quantitative trait loci (QTL) [3]. Others have functionally prioritized a single locus (*SNCA* [9] and *TMEM175* [10]) but these functional single locus experiments often do not scale up to loci with many genes. Other non-disease specific pipelines have been developed using epigenetic and chromatin conformation datasets in addition to expression QTL (eQTL) data [11,12], however some disease specific interpretation will be needed. Therefore, we have aggregated multiple datasets from several sources to create a versatile and user-friendly tool to prioritize specific genes and variants for additional PD GWAS and functional studies that aim to identify potential therapeutic targets. (Figure 1) https://pdgenetics.shinyapps.io/GWASBrowser/.

**Figure 1.**
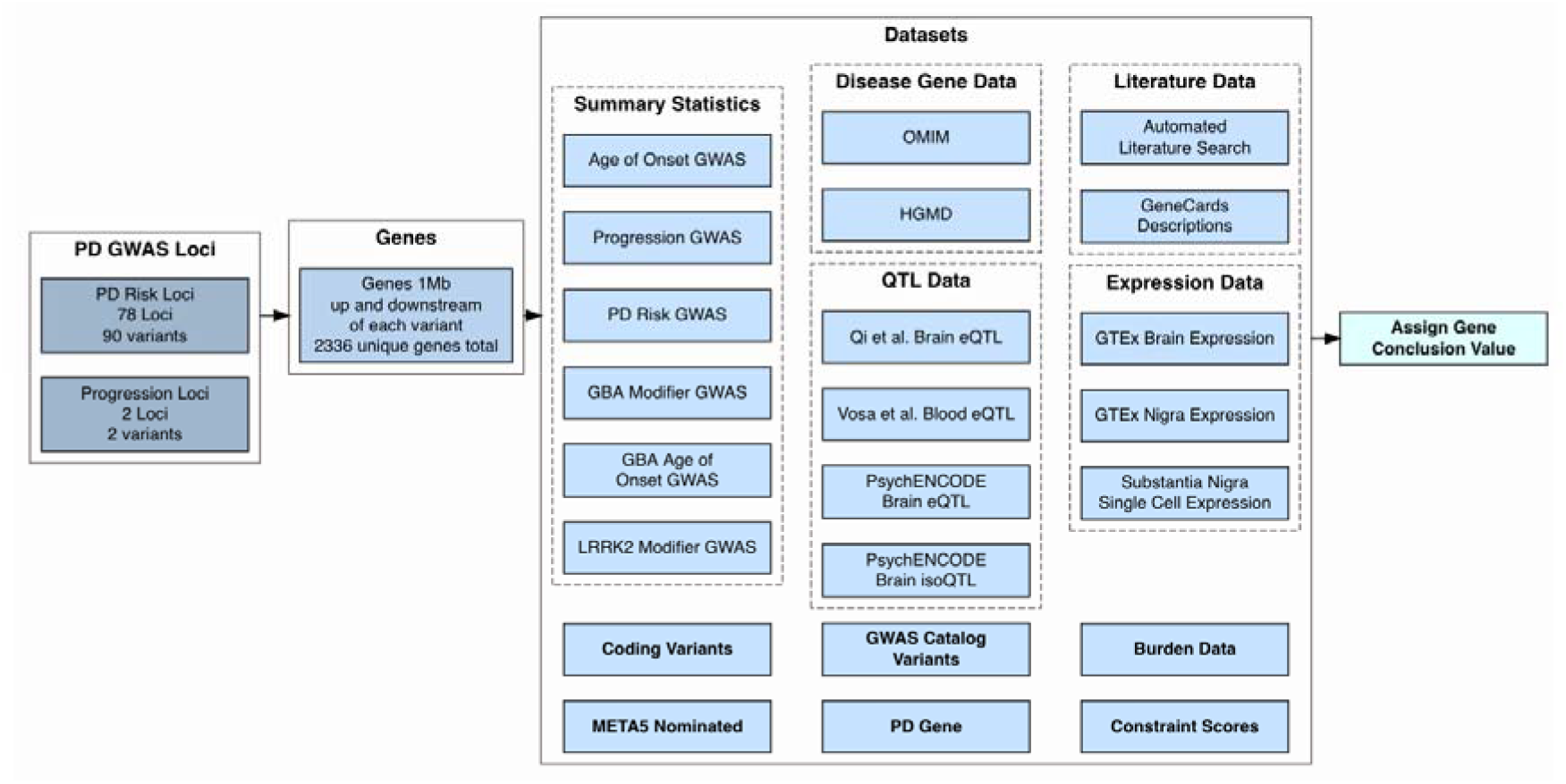
Flowchart of data gathered for the browser. Summary of the variants, genes and datasets included in the browser to prioritize genes for each locus. Datasets include genome wide association study (GWAS) summary statistics, known coding variants, META5 (Nalls et al 2019) nominated genes, online Mendelian inheritance in man (OMIM) and human gene mutation database (HGMD) disease genes, expression quantitative trait locus data (eQTL), variants from the GWAS catalog, known Parkinson’s disease genes, pubmed literature data, Genotype-Tissue Expression (GTEx) and single cell expression data, burden test data and constraint data.

## Methods

### GMAS loci and gene selection

All GWAS loci and summary statistics were gathered from our recent PD GWAS studies [2–4,13,14]. We selected all genes 1 Megabase (Mb) up and downstream of each significant variant from the hg19 reference genome [15]. This resulted in 2336 unique genes of interest. Genes in this 2Mb range were included in locus zoom plots created for each variant [16].

### Gene expression data

The Genotype-Tissue Expression (GTEx) portal was accessed on 02/12/2020 to obtain v8 gene expression data. Transcript per million (TPM) data was averaged across all available brain and substantia nigra (SN) samples individually for each gene. Average single cell RNA sequencing expression data was also included for SN astrocytes, SN dopaminergic neurons, SN endothelial cells, SN GABAergic cells, SN microglial cells, SN oligodendrocyte cells (ODC) and SN oligodendrocyte progenitor cells (OPC) (GSE140231, Agarwal et al 2020 under review). An arbitrary value of five TPM was chosen as the cut off point for significance in brain tissue, SN tissue and SN dopaminergic neuron averages. Genes with greater than five TPM in any of these three datasets were given a value of one in the evidence table in the “Brain Expression”, “Nigra Expression”, or “SN-Dop. Neuron Expression” column. Genes with no available expression data were set to “NA” in these columns.

### Expression quantitative trait loci data

Expression quantitative trait locus (eQTL) data was collected from the summary-data-based Mendelian randomization (SMR) website for brain tissues [17] and processed using SMR software tools [18]. Blood tissue eQTL data was collected from the eQTLGen consortium [19]. Additional brain tissue data was collected from the PsychENCODE project [20]. Locus compare plots were generated using GWAS data and the appropriate eQTL data to compare the distribution of eQTL and GWAS data [21]. Plots were omitted if they contained neither the locus risk variant nor a good proxy variant. Proxy variants were obtained using the LDlinkR library in R (https://www.r-project.org/) to find variants with r^2^ > 0.7 with the risk variant. Genes were given a value of one in the “QTL-brain” column in the evidence table if there was sufficient data to create brain locus compare plots using Qi et al. eQTL data, PsychENCODE eQTL data or PsychENCODE isoQTL data. Genes were given a value of one in the “QTL-blood” column in the evidence table if there was sufficient data in the blood tissue eQTL data to create a locus compare plot. Pearson correlation coefficients between GWAS and blood or brain eQTL/isoQTL p-values were calculated using R for each gene with sufficient data. The “QTL-correl” column in the evidence table was given a value of one if the magnitude of the correlation coefficient was greater than 0.3 in any of the gene’s locus compare plots. This column was given a value of “NA” if there were fewer than thirty datapoints in the locus compare plots for the gene.

### Literature search

Automated literature searches were performed in pubmed using the search term “[Gene Name][Title/Abstract], Parkinson’s[Title/Abstract]” and “[Gene Name][Title/Abstract]” to search for the respective gene and its occurence in PD literature using the rentrez package in R [22]. The number of search results were collected for all the genes in each locus to create a bar plot of pubmed hits for each locus. Genes with five or more search results in the PD and gene name search were given a score of one in the “Literature Search” column in the evidence table. A word cloud was generated for each gene using the PubMedWordcloud package in R [23]. Lastly, gene descriptions were obtained from GeneCards [24].

### Constraint data

Constraint data, a score describing how resistant to variation a gene is predicted to be, was downloaded from the gnomAD browser (https://gnomad.broadinstitute.org/downloads) [25]. Constraint z scores were included for synonymous variants and missense variants. A probability of being loss-of-function intolerant (pLI) score was included for loss of function variants. Observed/Expected variant values were included for these three variant types, along with the 90% confidence interval for each value.

### Burden data

Burden summary statistics were obtained from the most recent GWAS [3] and an exome sequencing study in PD (manuscript in process). In total, forty different burden tests on exome sequencing data and two different burden tests on imputed GWAS data were performed using only missense and loss-of-function variants using minor allele frequency cut-offs of 0.05 and 0.01. The minimum p-values of all forty exome burden tests were included for each gene in the burden table of the browser. The minimum p-value for the two burden tests on imputed data for each gene was included in the same table. These two minimum p-values were Bonferroni corrected by the number of genes with data in the burden test to determine significance (1466 genes for exome and 1014 genes for imputed). Genes with significant burden results in either exome or imputed data were given a value of one in the “Burden” column in the evidence table. Genes with no available burden data were given a value of “NA” in this column.

### Nalls et al 2019 Nominated Genes

The 2019 PD GWAS [3] used four QTL datasets to determine causal genes for each GWAS signal in the study. This data was obtained from Supplementary Table 1 of the Nalls et al 2019 GWAS paper. 70 of the 90 variants from this study were found to be associated with a putative causal gene. These genes were given a value of one in the “Nominated by META5” column of the evidence table.

### Parkinson’s disease and other disease genes

Genes known to be monogenic for PD or parkinsonism in the literature [1] were given a value of one in the “PD Gene” column in the evidence table. Disease gene data was gathered from the Human Gene Mutation Database [26] and OMIM [27]. For HGMD, only genes with variants classified as “DM” (disease causing mutations) were included. These genes were given a value of one in the “Disease Gene” column in the evidence table.

### Coding Variants

Coding variants in linkage disequilibrium (LD) with risk variants were obtained from internal databases. R^2^ and D’ LD scores were calculated using plink [28]. Combined annotation dependent depletion (CADD) scores were obtained from the CADD database using ANNOVAR [29]. Frequencies were obtained from the gnomAD database also using ANNOVAR [25,29].

### Associated Variant Phenotypes

Phenotypes of variants in LD with risk variants were obtained from the GWAS catalog v1.0.2 [30]. R^2^ and D’ LD scores were calculated using PLINK (v1.9) [31] from a large PD casecontrol reference set including over 40,000 individuals. Frequencies were obtained from the gnomAD database using ANNOVAR [25,29].

### Fine-Mapping

Variants in the PD meta-analysis summary statistics [3] were re-annotated to GRCh38p7 build positions using dbSNP build 151. If a variant’s dbSNP rs-id was not present in dbSNP build 151 it was excluded from further analysis. The summary stats were partitioned into risk regions or loci based on physical distance. Per chromosome these partitions were generated iteratively by finding the variant with the smallest P-value and extracting this variant and those variants within 1 Mb of it. The region of extracted proximal variants was checked against other extracted regions and if their edges were within 100 Kb the regions were merged. These iterations continued until no variant with a GWAS p-value less than 5e10^-8^ remained within the PD meta-analysis summary statistics.

The FINEMAP tool [3,32] was used to fine map these PD risk locus regions. Finemap uses Shotgun Stochastic Search [33] and Bayesian Model Averaging [34] to identify casual configurations of risk variants. The max number of causal variants within the configuration per locus was based on the number of independent risk signals detectable per locus. This number of independent risk signals per locus was estimated using the stepwise model selection procedure implemented in GCTA-COJO. Both the Finemap and GCTA-COJO [35,36] tools require linkage disequilibrium (LD) information for their search and modeling. For this purpose TOPMed freeze5b samples of European ancestry, available from dbGaP, were used as a LD reference panel; this panel included 16,257 samples.

### Browser Design

The GWAS locus browser is an R shiny application. Data was pre-compiled and loaded onto the application’s server and is not obtained from external databases in real time. The GWAS locus browser is an open source project. The code is available on our github https://github.com/neurogenetics/GWAS_locus_browser.

## Results

### Data-browser

The PD GWAS locus browser (https://pdgenetics.shinyapps.io/GWASBrowser/) is an online platform to assist researchers with the prioritization of genes located within PD GWAS loci. It includes multiple layers of data, including: GWAS statistic, eQTL, burden, expression, constraint, and literature data, and a flexible scoring system that users may configure for their own needs. In total we included 92 GWAS variants located on 80 GWAS loci and a total of 2336 genes. Interestingly, several PD GWAS hits show high correlation and overlap (R^2^>0.8 and D’>0.9) (Supplementary Table 1) with other disease risk signals including disease such as inflammatory bowel disease (Locus 2) [37,38], neuroticism (Loci 29, 62, 69 and 73) [39–41], body mass index (Loci 4, 11, 61) [42,43] and insomnia (Locus 73) [44]. Below we describe two use-case scenarios on how this application could be used for prioritizing genes from PD GWAS loci.

### Use case scenario 1, locus 16, rs11707416

A possible use case for the browser exists on locus 16 for the risk variant rs11707416. This variant was discovered in the most recent PD GWAS (P=1.13E-10, OR=0.94, SE=0.0097) [3] and the closest gene is *MED12L*. The locus zoom plot shows a clear GWAS signal for this region (Figure 2A) however, no genes in this locus were prioritized using current methods in the PD GWAS. Using the PD GWAS browser default settings, *P2RY12* (Purinergic Receptor P2Y) has the highest conclusion score (7) in the evidence per gene table, meaning it has the highest sum of dataset scores for that locus. Locus compare eQTL plots show some correlation in both brain and blood, indicating there is some overlap in the distribution of eQTL and GWAS data for this gene (Figure 2B). There is one common coding variant at this locus, but it is located within *MED12L* (Mediator complex subunit 12-like; NM_053002:exon25:c.G3629A:p.R1210Q) and not *P2RY12. MED12L* has a lower conclusion score (3) than *P2RY12*, suggesting that even with this coding variant, *MED12L* is not the primary candidate in this locus. This is a very complex and unusual locus, in that there are multiple genes encoded within an intron of an isoform of *MED12L* (*P2RY12, P2RY13, P2RY14, GPR171*), and even more interesting is that the missense variant which changes an amino acid of *MED12L* is located in an intron of the much smaller gene *P2RY12* (Figure 2A). Despite the uniqueness of the locus structure, we will focus on *P2RY12* as the primary candidate nominated by the PD GWAS browser datasets.

**Figure 2.**
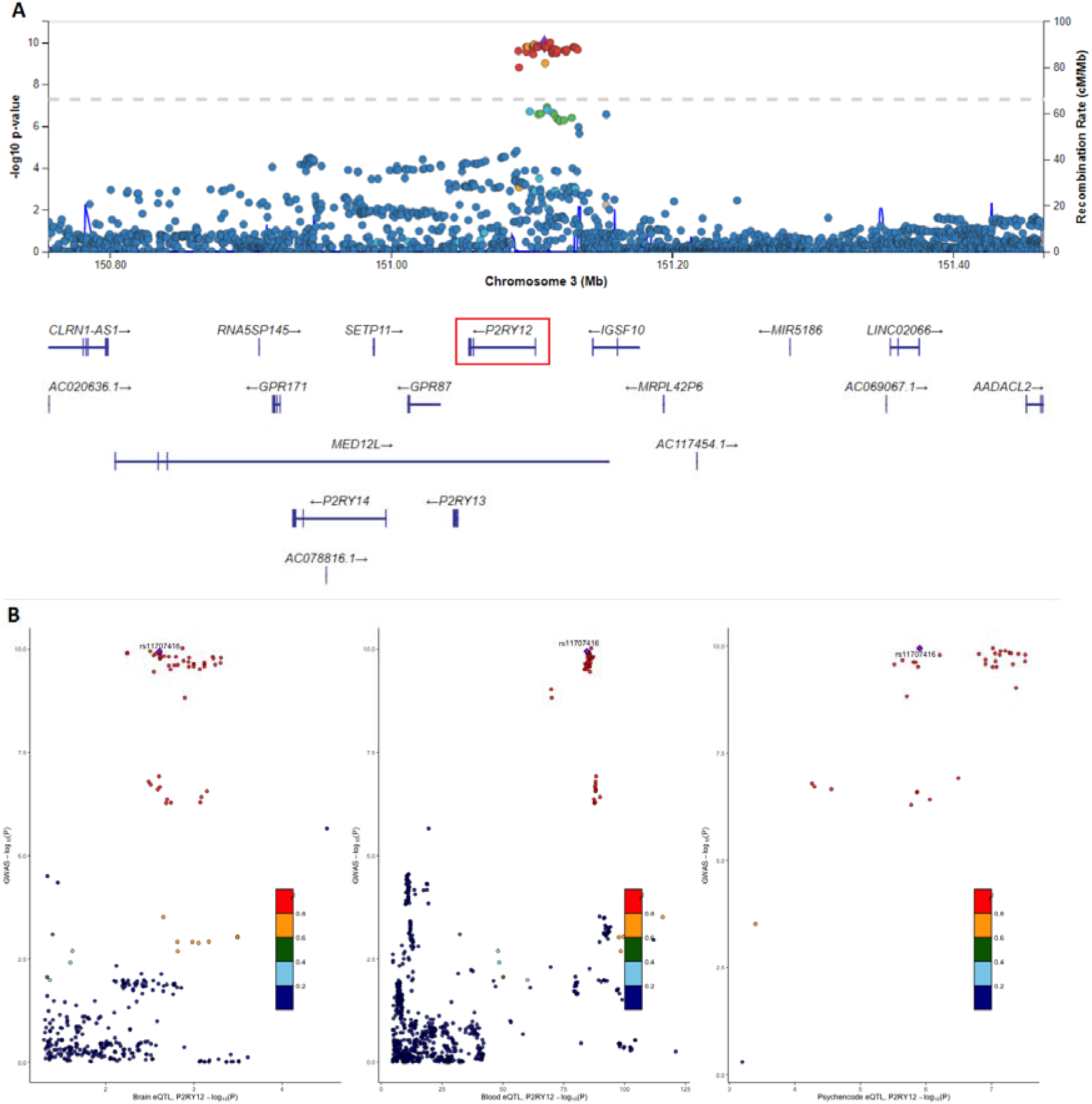
Locus zoom and locus compare plots for locus 16 and variant rs11707416 and *P2RY12*. (A) The locus zoom Manhattan plot for risk variant rs11707416 on locus 16. The risk variant is uniquely colored purple and all other variants are colored by their r^2^ value. Recombination rate peaks are plotted in blue. Nearby genes are included at the bottom, with *P2RY12* highlighted. (B) Locus compare plots for *P2RY12* on locus 16 plot −log_10_P-values for Nalls et al 2019 GWAS data (y-axis) and for different eQTL datasets (x-axis). Datasets available for *P2RY12* are Qi et al. brain eQTL, Vosa et al. blood eQTL and PsychENCODE brain eQTL (left to right). Variants are colored by their r^2^ value and the risk variant is labelled and uniquely colored purple.

Expression of *P2RY12* was significant in all included databases (GTEx brain, GTEx SN and single cell dopaminergic neuron data). However, it appears that *P2RY12* has much higher expression in astrocytes and microglia than neurons (Figure 3). The disease gene section shows that *P2RY12* has been linked to platelet type-8 bleeding disorder [45]. This disorder appears to be caused by dominant negative mutations in the *P2RY12* gene that disrupt the homo-dimerization of the receptor which is required for normal function [46]. No direct links between *P2RY12* and PD have been reported in the literature, but *P2RY12* is a widely studied gene with roles suggested in neuroinflammation, apoptosis and autophagy, pathways that are relevant to PD and other neurodegenerative diseases [47–49]. Experiments have been done to characterize expression patterns of *P2RY12* in microglia and its role in neuroinflammation [50]. As these experiments focused on Alzheimer’s disease, it would be useful to build upon them in the PD context in the future. The possible role of *P2RY12 in PD* should be further analyzed using functional tests in cell and animal models.

**Figure 3.**
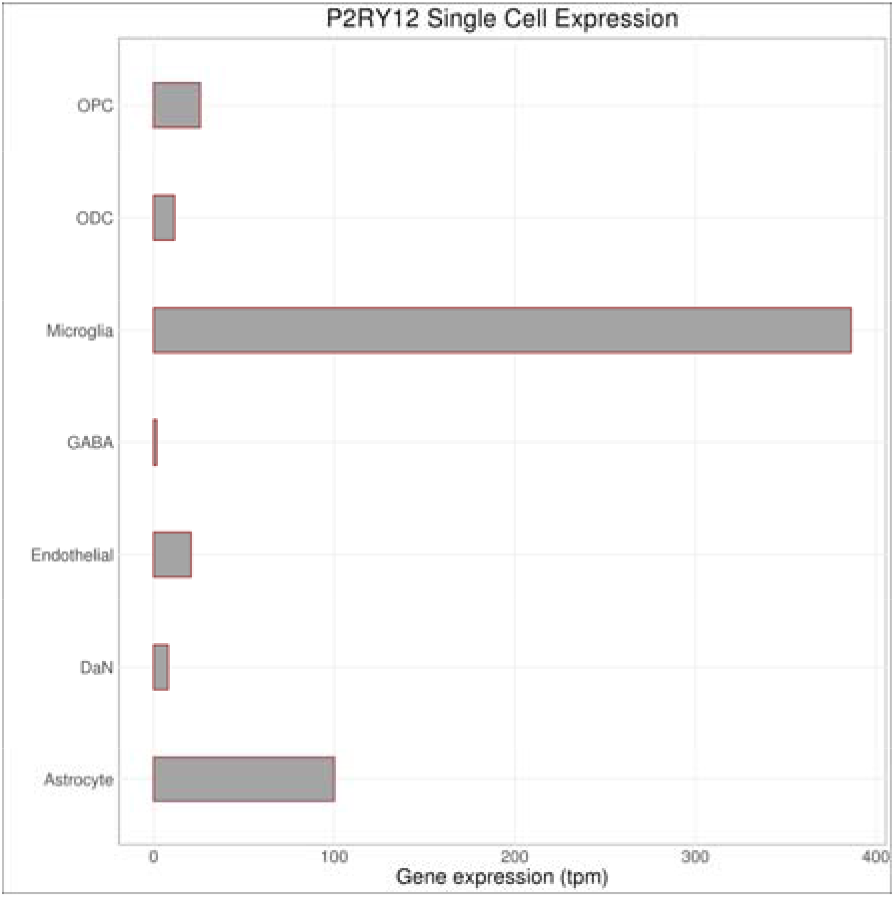
Bar plot for single cell expression of *P2RY12* on locus 16. Transcript per million (TPM) data for *P2RY12* was averaged across all samples for seven different cell types from the substantia nigra. These include oligodendrocyte progenitor cells (OPC), oligodendrocyte cells (ODC), microglia, GABAergic neurons (GABA), endothelial cells, dopaminergic neurons (DaN) and astrocytes.

### Use case scenario 2: Locus 78, rs2248244

We have chosen a second use case for the browser at locus 78, for the risk variant rs2248244. This variant was discovered in the most recent PD GWAS (P=2.74E-11, OR=1.074, SE=0.0107) and its nearest gene, *DYRK1A*, was nominated as the causal gene in that study. The locus zoom plot shows a clear GWAS signal for this region (Figure 4A). The default settings in the evidence per gene table nominate *DYRK1A* as the top candidate with a conclusion score of 10, which is higher than any of other genes within the locus (second highest is 4).

**Figure 4.**
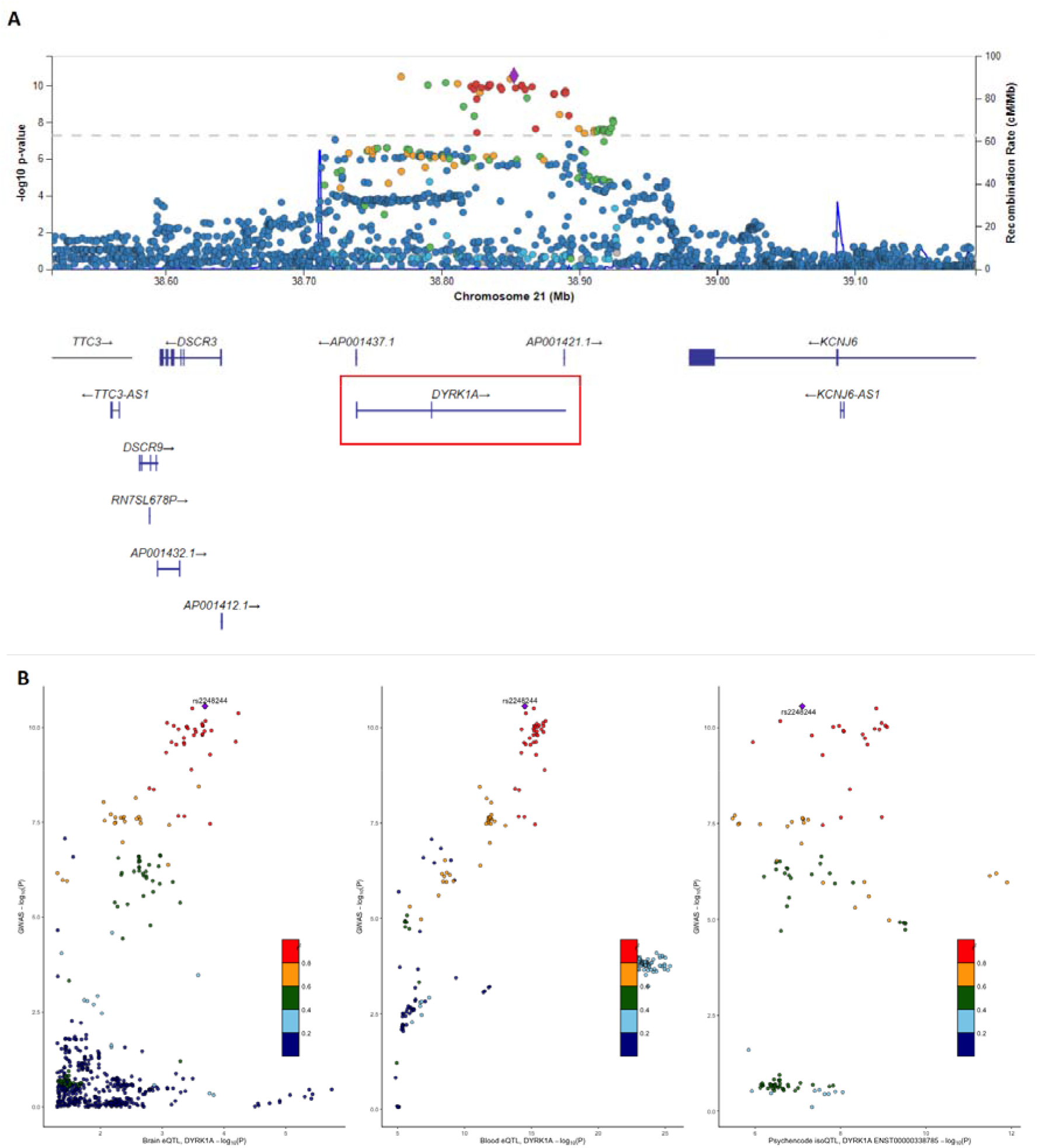
Locus zoom and locus compare plots for locus 78 and variant rs2248244 and *DYRK1A*. (A) The locus zoom Manhattan plot for showing risk variant rs2248244 at locus 78. The risk variant is uniquely colored purple and all other variants are colored by their r^2^ value. Recombination rate peaks are plotted in blue. Nearby genes are included at the bottom, with *DYRK1A* highlighted. (B) Locus compare plots for *DYRK1A* on locus 78 plot - log10P-values for Nalls et al 2019 GWAS data (y-axis) and for different eQTL datasets (x-axis). Datasets available for *DYRK1A* are Qi et al. brain eQTL, Vosa et al. blood eQTL and PsychENCODE brain isoQTL (left to right). Variants are colored by their r^2^ value and the risk variant is labelled and uniquely colored purple.

The brain and blood eQTL plots and the isoQTL plot show good correlation between GWAS and QTL values (Figure 4B). No coding variants or other known associated disease variants exist for this locus. *DYRK1A* showed significant gene expression in all databases (GTEx brain, GTEx SN and single cell dopaminergic neuron data) (Figure 5). Constraint data for *DYRK1A* shows a low 90% CI for loss of function variation and a pLI of 1, suggesting significant intolerance to loss of function variation. However, burden test results show no significant change in variants for *DYRK1A* after Bonferroni correction. FINEMAP results of this locus nominated several variants with rs2248244 and rs11701722 both intronic with the highest probability score.

**Figure 5.**
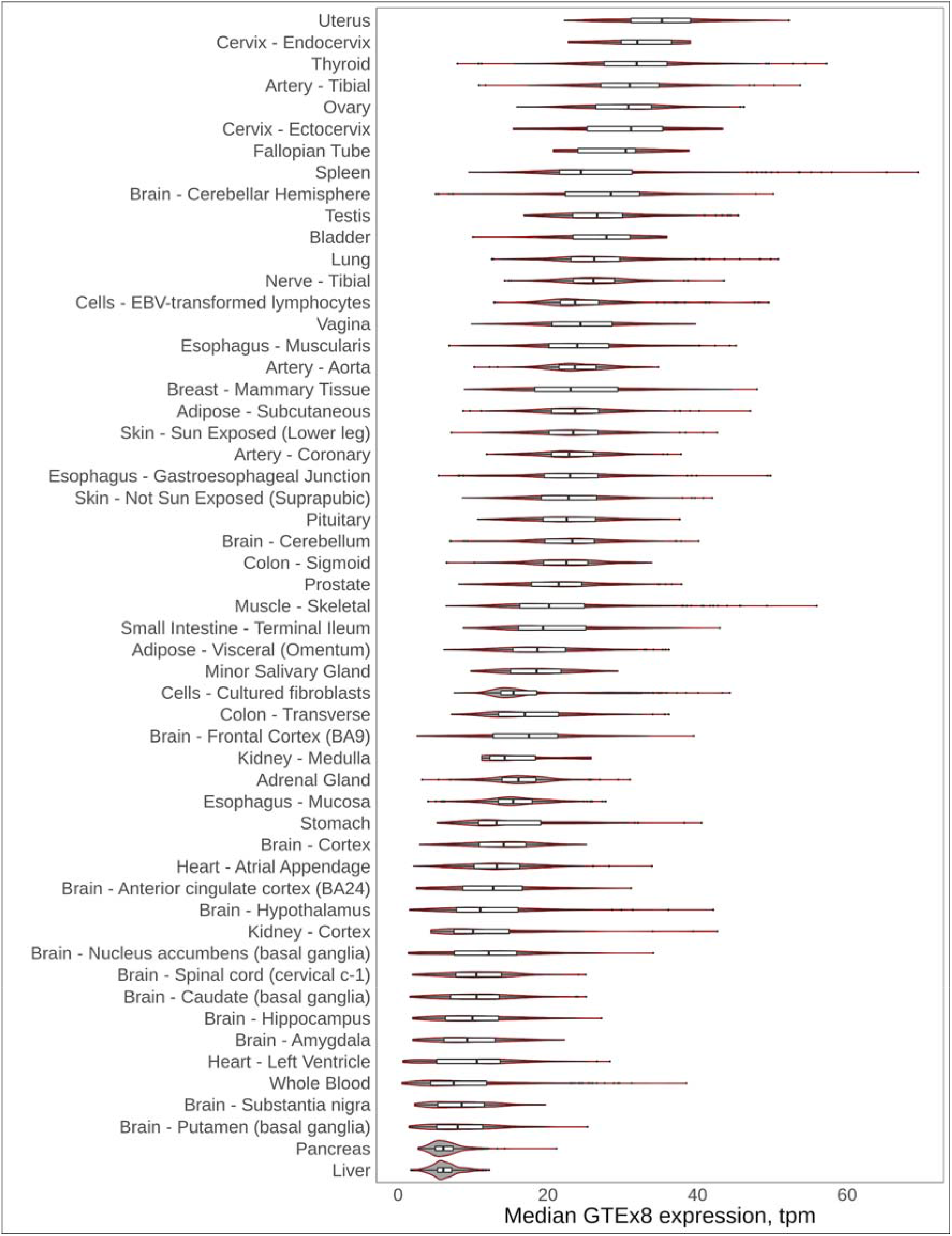
GTEx violin plot for *DYRK1A* on locus 78. *DYRK1A* transcript per million (TPM) data was averaged across all available samples from GTEx v8 data for different tissues. Distribution and probability of expression level is included for each tissue.

*DYRK1A* is located in the Down Syndrome Critical Region (DSCR) on chromosome 21 and various deletions and single nucleotide variants have been linked to autosomal dominant mental retardation-7 (MRD7) [51–53]. This data is not enough to conclude that *DYRK1A* is the relevant risk gene at this locus, however, *DYRK1A* also has a significant representation in PD literature. Previous studies have suggested that *DYRK1A* encodes a kinase that can phosphorylate α-synuclein and Pink1 in mammalian cells [54–57]. Studies in mouse models indicate that haploinsufficiency of *DYRK1A* leads to a reduction in dopaminergic neurons while increasing the dosage results in more dopaminergic neurons by altering apoptosis [54–56]. All of these studies combined indicate that loss of *DYRK1A* expression may influence the number of dopaminergic neurons and thereby the development of PD. Therefore, the PD GWAS browser predicts *DYRK1A* is a good candidate for more functional PD studies in cells and mouse models as well as increasing individuals in human genetic studies to characterize the molecular mechanism underlying risk at this locus.

## Discussion

GWAS have identified numerous risk loci for a large number of diseases (https://www.ebi.ac.uk/gwas/) [30]. The current bottleneck after completing these studies is identifying the causal genes and variants underlying the GWAS signals. There is a large discrepancy between the number of new GWAS loci identified and the number of studies that molecularly characterize these loci to identify a causative variant or gene. Currently, the number of PD loci that are functionally validated is very low and mostly includes genes that are known to cause monogenic forms of PD. GWAS has identified a number of these pleiotropic genes at various loci such as *SNCA* (locus 23), *GBA* (locus1), *LRRK2* (locus 49) and *VPS13C* (locus 59). Current evidence suggests that *TMEM175* (locus 19 [10]), *CTSB* (locus 37 [8,13]) and *GCH1* (locus 56, [58]) are also pleiotropic and lead to PD by multiple mechanisms.

Our goal is to provide the PD research community with a tool that catalogs all significant PD GWAS signals and helps prioritize the genes at each locus for functional studies. The overall significance score for each gene is displayed in the “Conclusion” column of the evidence table. This score is the sum of all other numerical values from each dataset for the respective gene. The inclusion of genes 1 Mb up and downstream of each variant is an arbitrary cutoff and may limit the ability of the browser to accurately prioritize genes. This limitation will come into effect when variants affect genes outside this range, however prior studies have found that functional noncoding variants are often located within this 2Mb window. Additionally, some loci contain more than one significant variant (eg locus 1, *GBA* with three reported independent signals) and until a way to detect precise causative variants is developed, it is assumed that all significant variants within the same locus will impact the same gene.

While all of the datasets included in the PD browser may contribute to the prioritization of PD genes, each dataset comes with its own limitations that should be taken into account when considering the conclusion value. Blood and brain eQTL data was included to identify genes with similar GWAS and eQTL data distribution. While similar distributions may suggest causality, the power of these datasets may reduce their importance. Blood eQTL data will have more statistical power than brain data due to its larger sample size, but it may be less relevant to PD. Another important note is that we just test for cis eQTLs, since we simply do not have power to detect robust trans-QTLs. Scoring for the “QTL-brain” and “QTL-blood” column is not indicative of the causality of a gene because it relies on the existence of eQTL data, not the information provided by the data. While the “QTL-correl” column gives more insight into the eQTL data, the Pearson correlation coefficient cutoff significance value of - 0.3 and 0.3 are quite broad. This arbitrary value was chosen to account for the low number of genes that qualified for locus compare plots. Additionally, eQTL data is not available for all genes of interest. For these reasons, the correlation between eQTL and GWAS data does not guarantee causality for a gene, but is still good evidence for prioritization. Gene expression data was included to account for possible increased expression of genes associated with GWAS variants in relevant cell types and tissues. GTEx Transcript per million (TPM) v8 data was used to measure gene expression. We focused on brain tissue, SN tissue, and SN dopaminergic neuron data because of their established role in PD and enrichment in PD GWAS loci [3]. However, genes do not necessarily need to be expressed in the substantia nigra or other brain tissues and cell types to increase the risk of PD. Constraint data was included to identify genes that are intolerant to specific types of variation, suggesting conservation. Therefore, we suspect genes with significant intolerance may be causal because normal variation in that gene is unlikely. We used a significance cutoff of 0.35 for the upper limit of the 90% confidence interval of the observed/expected value as suggested by gnomAD [25]. It is possible that variations within genes may not be associated with PD, suggesting that low intolerance/constraint scores aren’t necessarily causal. Previously published burden test results were included to account for genes thought to have a significant burden of rare variants in PD (Nalls et al., 2019). However, this does not guarantee causality, because causal genes may be tagged by common variants instead. The literature count was included to quickly measure the significance of genes in previously published research. However, existing studies can easily be biased, and our automated search of the literature does not account for this. Additionally, some genes are difficult to identify in automated literature searches due to nomenclature changing over time or their similarity to common names. Examples of these name complications include *SHE* and *MAL* (sometimes used for “mean axonal length” instead of the gene encoding “Myelin And Lymphocyte Protein”).

Overall, for approximately half of all PD GWAS loci, an educated guess can be made based on the data included in our browser to choose the most likely candidate gene underlying the GWAS risk signal. For some loci where there is no obvious candidate (eg loci 34, 50 and 74) but our browser may still help prioritize candidate genes with the help of additional specific reference data. It should also be noted that there is not a “one-fits all” efficient scoring system for the prioritization of genes from under GWAS peaks. Additionally, each included dataset has clear limitations as discussed above. Therefore we included a weight option for each data column in the evidence table. This will allow users to assign weights (0-4, 0 for no points and 4 for 4 points) to different columns to alter the significance of the data in the final conclusion summation based on the datasets they deem most important.

In summary, we present here an online platform that allows for prioritization of genes within PD GWAS loci. We highlight two examples (*P2RY12* and *DYRK1A*) but many other interesting gene-candidates can be identified using this application. The platform is designed to be versatile, flexible and easily expandable when more loci or datasets of interest become available. By using this platform, GWAS follow-up studies can systematically prioritize genes based on publicly available datasets which may help improve the design of functional experiments. In turn, this workflow could help nominate these genes as potential therapeutic targets worthy of translating to the clinic.

## Acknowledgements and Funding

We would also like to thank all members of the International Parkinson Disease Genomics Consortium (IPDGC). For a complete overview of members, acknowledgements and funding, please see the Supplemental data and/or http://pdgenetics.org/partners. This work was supported in part by the Intramural Research Programs of the National Institute of Neurological Disorders and Stroke (NINDS), the National Institute on Aging (NIA), and the National Institute of Environmental Health Sciences both part of the National Institutes of Health, Department of Health and Human Services; project numbers 1ZIA-NS003154, Z01-AG000949-02 and Z01-ES101986. We thank the research participants and employees of 23andMe for making this work possible. CW is supported by the UK Dementia Research Institute funded by the Medical Research Council (MRC), Alzheimer’s Society and Alzheimer’s Research UK. CS is supported by the Ser Cymru II programme which is part-funded by Cardiff University and the European Regional Development Fund through the Welsh Government. Data were generated as part of the PsychENCODE Consortium supported by: U01MH103339, U01MH103365, U01MH103392, U01MH103340, U01MH103346, R01MH105472, R01MH094714, R01MH105898, R21MH102791, R21MH105881, R21MH103877, and P50MH106934 awarded to: Schahram Akbarian (Icahn School of Medicine at Mount Sinai), Gregory Crawford (Duke), Stella Dracheva (Icahn School of Medicine at Mount Sinai), Peggy Farnham (USC), Mark Gerstein (Yale), Daniel Geschwind (UCLA), Thomas M. Hyde (LIBD), Andrew Jaffe (LIBD), James A. Knowles (USC), Chunyu Liu (UIC), Dalila Pinto (Icahn School of Medicine at Mount Sinai), Nenad Sestan (Yale), Pamela Sklar (Icahn School of Medicine at Mount Sinai), Matthew State (UCSF), Patrick Sullivan (UNC), Flora Vaccarino (Yale), Sherman Weissman (Yale), Kevin White (UChicago) and Peter Zandi (JHU). The Genotype-Tissue Expression (GTEx) Project was supported by the Common Fund of the Office of the Director of the National Institutes of Health, and by NCI, NHGRI, NHLBI, NIDA, NIMH, and NINDS. The data used for the analyses described in this manuscript were obtained from the GTEx Portal on 02/12/2020. Molecular data for the Trans-Omics in Precision Medicine (TOPMed) program was supported by the National Heart, Lung and Blood Institute (NHLBI). Genome sequencing for “NHLBI TOPMed: Whole Genome Sequencing and Related Phenotypes in the Framingham Heart Study” (phs000974.v1.p1) was performed at the Broad Institute of MIT and Harvard (HHSN268201500014C). RNASeq for “NHLBI TOPMed: Whole Genome Sequencing and Related Phenotypes in the Framingham Heart Study” (phs000974.v1.p1)” was performed at the University of Washington Northwest Genomics Center (HHSN268201600032I). Genome sequencing for “NHLBI TOPMed: The Jackson Heart Study” (phs000964.v1.p1) was performed at the University of Washington Northwest Genomics Center (HHSN268201100037C). Core support including centralized genomic read mapping and genotype calling, along with variant quality metrics and filtering were provided by the TOPMed Informatics Research Center (3R01HL-117626-02S1; contract HHSN268201800002I). Core support including phenotype harmonization, data management, sample-identity QC, and general program coordination were provided by the TOPMed Data Coordinating Center (R01HL-120393; U01HL-120393; contract HHSN268201800001I). We gratefully acknowledge the studies and participants who provided biological samples and data for TOPMed.

## Author contributions

Design and concept: FPG, JJK, MBM, CB, AS

App development: FPG, JJK, MBM

Additional data contribution and analysis: All

Manuscript drafting: FPG, XR, CB

Manuscript revision: All

## References

1. Blauwendraat C, Nalls MA, Singleton AB. The genetic architecture of Parkinson’s disease. Lancet Neurol. 2020;19: 170–178.

2. Iwaki H, Blauwendraat C, Leonard HL, Kim JJ, Liu G, Maple-Grødem J, et al. Genomewide association study of Parkinson’s disease clinical biomarkers in 12 longitudinal patients’ cohorts. Mov Disord. 2019. doi:10.1002/mds.27845

3. Nalls MA, Blauwendraat C, Vallerga CL, Heilbron K, Bandres-Ciga S, Chang D, et al. Identification of novel risk loci, causal insights, and heritable risk for Parkinson’s disease: a meta-analysis of genome-wide association studies. Lancet Neurol. 2019;18: 1091–1102.

4. Blauwendraat C, Heilbron K, Vallerga CL, Bandres-Ciga S, von Coelln R, Pihlstrøm L, et al. Parkinson’s disease age at onset genome-wide association study: Defining heritability, genetic loci, and α-synuclein mechanisms. Mov Disord. 2019;34: 866–875.

5. Lettice LA, Heaney SJH, Purdie LA, Li L, de Beer P, Oostra BA, et al. A long-range Shh enhancer regulates expression in the developing limb and fin and is associated with preaxial polydactyly. Hum Mol Genet. 2003;12: 1725–1735.

6. Claussnitzer M, Dankel SN, Kim K-H, Quon G, Meuleman W, Haugen C, et al. FTO Obesity Variant Circuitry and Adipocyte Browning in Humans. N Engl J Med. 2015;373: 895–907.

7. Hook PW, McClymont SA, Cannon GH, Law WD, Morton AJ, Goff LA, et al. Single-Cell RNA-Seq of Mouse Dopaminergic Neurons Informs Candidate Gene Selection for Sporadic Parkinson Disease. Am J Hum Genet. 2018;102: 427–446.

8. Li YI, Wong G, Humphrey J, Raj T. Prioritizing Parkinson’s disease genes using population-scale transcriptomic data. Nat Commun. 2019;10: 994.

9. Soldner F, Stelzer Y, Shivalila CS, Abraham BJ, Latourelle JC, Barrasa MI, et al. Parkinson-associated risk variant in distal enhancer of α-synuclein modulates target gene expression. Nature. 2016;533: 95–99.

10. Jinn S, Blauwendraat C, Toolan D, Gretzula CA, Drolet RE, Smith S, et al. Functionalization of the TMEM175 p.M393T variant as a risk factor for Parkinson disease. Hum Mol Genet. 2019;28: 3244–3254.

11. Peat G, Jones W, Nuhn M, Marugán JC, Newell W, Dunham I, et al. The Open Targets Post-GWAS analysis pipeline. Bioinformatics. 2020. doi:10.1093/bioinformatics/btaa020

12. Watanabe K, Taskesen E, van Bochoven A, Posthuma D. Functional mapping and annotation of genetic associations with FUMA. Nat Commun. 2017;8: 1826.

13. Blauwendraat C, Reed X, Krohn L, Heilbron K, Bandres-Ciga S, Tan M, et al. Genetic modifiers of risk and age at onset in GBA associated Parkinson’s disease and Lewy body dementia. Brain. 2020;143: 234–248.

14. Iwaki H, Blauwendraat C, Makarious MB, Bandrés-Ciga S, Leonard HL, Gibbs JR, et al. Penetrance of Parkinson’s Disease in LRRK2 p.G2019S Carriers Is Modified by a Polygenic Risk Score. Mov Disord. 2020. doi:10.1002/mds.27974

15. Haeussler M, Zweig AS, Tyner C, Speir ML, Rosenbloom KR, Raney BJ, et al. The UCSC Genome Browser database: 2019 update. Nucleic Acids Res. 2019;47: D853–D858.

16. Pruim RJ, Welch RP, Sanna S, Teslovich TM, Chines PS, Gliedt TP, et al. LocusZoom: regional visualization of genome-wide association scan results. Bioinformatics. 2010;26: 2336–2337.

17. Qi T, Wu Y, Zeng J, Zhang F, Xue A, Jiang L, et al. Identifying gene targets for brain-related traits using transcriptomic and methylomic data from blood. Nat Commun. 2018;9: 2282.

18. Zhu Z, Zhang F, Hu H, Bakshi A, Robinson MR, Powell JE, et al. Integration of summary data from GWAS and eQTL studies predicts complex trait gene targets. Nat Genet. 2016;48: 481–487.

19. Võsa U. Unraveling the polygenic architecture of complex traits using blood eQTL metaanalysis. bioRxiv. 2018. doi:10.1101/447367

20. Wang D, Liu S, Warrell J, Won H, Shi X, Navarro FCP, et al. Comprehensive functional genomic resource and integrative model for the human brain. Science. 2018;362. doi:10.1126/science.aat8464

21. Liu B, Gloudemans M, Montgomery S. LocusCompare: A Tool to Visualize Pairs of Association. 2018. Available: http://locuscompare.com/

22. Winter D. rentrez: An R package for the NCBI eUtils API. The R Journal. 2017. pp. 520–526. doi:10.32614/rj-2017-058

23. Fan Y. Pubmedwordcloud. 2014. Available: http://felixfan.github.io/PubMedWordcloud/

24. Stelzer G, Rosen N, Plaschkes I, Zimmerman S, Twik M, Fishilevich S, et al. The GeneCards Suite: From Gene Data Mining to Disease Genome Sequence Analyses. Curr Protoc Bioinformatics. 2016;54: 1.30.1–1.30.33.

25. Karczewski KJ, Francioli LC, Tiao G, Cummings BB, Alföldi J, Wang Q, et al. Variation across 141,456 human exomes and genomes reveals the spectrum of loss-of-function intolerance across human protein-coding genes. bioRxiv. 2019. p. 531210. doi:10.1101/531210

26. Stenson PD, Mort M, Ball EV, Evans K, Hayden M, Heywood S, et al. The Human Gene Mutation Database: towards a comprehensive repository of inherited mutation data for medical research, genetic diagnosis and next-generation sequencing studies. Hum Genet. 2017;136: 665–677.

27. McKusick-Nathans Institute of Genetic Medicine, Johns Hopkins University (Baltimore, MD). Online Mendelian Inheritance in Man, OMIM®. [cited 15 Feb 2020]. Available: https://omim.org/

28. Purcell S, Neale B, Todd-Brown K, Thomas L, Ferreira MAR, Bender D, et al. PLINK: a tool set for whole-genome association and population-based linkage analyses. Am J Hum Genet. 2007;81: 559–575.

29. Wang K, Li M, Hakonarson H. ANNOVAR: functional annotation of genetic variants from high-throughput sequencing data. Nucleic Acids Res. 2010;38: e164.

30. Buniello A, MacArthur JAL, Cerezo M, Harris LW, Hayhurst J, Malangone C, et al. The NHGRI-EBI GWAS Catalog of published genome-wide association studies, targeted arrays and summary statistics 2019. Nucleic Acids Res. 2019;47: D1005–D1012.

31. Chang CC, Chow CC, Tellier LC, Vattikuti S, Purcell SM, Lee JJ. Second-generation PLINK: rising to the challenge of larger and richer datasets. Gigascience. 2015;4: 7.

32. Benner C, Havulinna AS, Salomaa V, Ripatti S, Pirinen M. Refining fine-mapping: effect sizes and regional heritability. Genetics. bioRxiv; 2018. pp. R111–9.

33. Hans C, Dobra A, West M. Shotgun Stochastic Search for “Largep” Regression. Journal of the American Statistical Association. 2007. pp. 507–516. doi:10.1198/016214507000000121

34. Kass RE, Raftery AE. Bayes Factors. Journal of the American Statistical Association. 1995. pp. 773–795. doi:10.1080/01621459.1995.10476572

35. Yang J, Genetic Investigation of ANthropometric Traits (GIANT) Consortium, Ferreira T, Morris AP, Medland SE, Madden PAF, et al. Conditional and joint multiple-SNP analysis of GWAS summary statistics identifies additional variants influencing complex traits. Nature Genetics. 2012. pp. 369–375. doi:10.1038/ng.2213

36. Yang J, Lee SH, Goddard ME, Visscher PM. GCTA: a tool for genome-wide complex trait analysis. Am J Hum Genet. 2011;88: 76–82.

37. de Lange KM, Moutsianas L, Lee JC, Lamb CA, Luo Y, Kennedy NA, et al. Genomewide association study implicates immune activation of multiple integrin genes in inflammatory bowel disease. Nat Genet. 2017;49: 256–261.

38. Jostins L, Ripke S, Weersma RK, Duerr RH, McGovern DP, Hui KY, et al. Host-microbe interactions have shaped the genetic architecture of inflammatory bowel disease. Nature. 2012;491: 119–124.

39. Luciano M, Hagenaars SP, Davies G, Hill WD, Clarke T-K, Shirali M, et al. Association analysis in over 329,000 individuals identifies 116 independent variants influencing neuroticism. Nat Genet. 2018;50: 6–11.

40. Nagel M, Watanabe K, Stringer S, Posthuma D, van der Sluis S. Item-level analyses reveal genetic heterogeneity in neuroticism. Nat Commun. 2018;9: 905.

41. Baselmans BML, Jansen R, Ip HF, van Dongen J, Abdellaoui A, van de Weijer MP, et al. Multivariate genome-wide analyses of the well-being spectrum. Nat Genet. 2019;51: 445–451.

42. Wang H, Zhang F, Zeng J, Wu Y, Kemper KE, Xue A, et al. Genotype-by-environment interactions inferred from genetic effects on phenotypic variability in the UK Biobank. Sci Adv. 2019;5: eaaw3538.

43. Akiyama M, Okada Y, Kanai M, Takahashi A, Momozawa Y, Ikeda M, et al. Genomewide association study identifies 112 new loci for body mass index in the Japanese population. Nat Genet. 2017;49: 1458–1467.

44. Jansen PR, Watanabe K, Stringer S, Skene N, Bryois J, Hammerschlag AR, et al. Genome-wide analysis of insomnia in 1,331,010 individuals identifies new risk loci and functional pathways. Nat Genet. 2019;51: 394–403.

45. Cattaneo M, Zighetti ML, Lombardi R, Martinez C, Lecchi A, Conley PB, et al. Molecular bases of defective signal transduction in the platelet P2Y12 receptor of a patient with congenital bleeding. Proc Natl Acad Sci U S A. 2003;100: 1978–1983.

46. Mundell SJ, Rabbolini D, Gabrielli S, Chen Q, Aungraheeta R, Hutchinson JL, et al. Receptor homodimerization plays a critical role in a novel dominant negative P2RY12 variant identified in a family with severe bleeding. J Thromb Haemost. 2018;16: 44–53.

47. Pi S, Mao L, Chen J, Shi H, Liu Y, Guo X, et al. The P2RY12 receptor promotes VSMC-derived foam cell formation by inhibiting autophagy in advanced atherosclerosis. Autophagy. 2020. doi:10.1080/15548627.2020.1741202

48. Blume ZI, Lambert JM, Lovel AG, Mitchell DM. Microglia in the developing retina couple phagocytosis with the progression of apoptosis via P2RY12 signaling. Dev Dyn. 2020. doi:10.1002/dvdy.163

49. van Wageningen TA, Vlaar E, Kooij G, Jongenelen CAM, Geurts JJG, van Dam A-M. Regulation of microglial TMEM119 and P2RY12 immunoreactivity in multiple sclerosis white and grey matter lesions is dependent on their inflammatory environment. Acta Neuropathol Commun. 2019;7: 206.

50. Walker DG, Tang TM, Mendsaikhan A, Tooyama I, Serrano GE, Sue LI, et al. Patterns of Expression of Purinergic Receptor P2RY12, a Putative Marker for Non-Activated Microglia, in Aged and Alzheimer’s Disease Brains. International Journal of Molecular Sciences. 2020. p. 678. doi:10.3390/ijms21020678

51. van Bon BWM, Hoischen A, Hehir-Kwa J, de Brouwer APM, Ruivenkamp C, Gijsbers ACJ, et al. Intragenic deletion in DYRK1A leads to mental retardation and primary microcephaly. Clin Genet. 2011;79: 296–299.

52. Courcet J-B, Faivre L, Malzac P, Masurel-Paulet A, Lopez E, Callier P, et al. The DYRK1A gene is a cause of syndromic intellectual disability with severe microcephaly and epilepsy. J Med Genet. 2012;49: 731–736.

53. O’Roak BJ, Vives L, Fu W, Egertson JD, Stanaway IB, Phelps IG, et al. Multiplex targeted sequencing identifies recurrently mutated genes in autism spectrum disorders. Science. 2012;338: 1619–1622.

54. Chiu C-C, Yeh T-H, Chen R-S, Chen H-C, Huang Y-Z, Weng Y-H, et al. Upregulated Expression of MicroRNA-204-5p Leads to the Death of Dopaminergic Cells by Targeting DYRK1A-Mediated Apoptotic Signaling Cascade. Front Cell Neurosci. 2019;13: 399.

55. Barallobre MJ, Perier C, Bové J, Laguna A, Delabar JM, Vila M, et al. DYRK1A promotes dopaminergic neuron survival in the developing brain and in a mouse model of Parkinson’s disease. Cell Death & Disease. 2014. pp. e1289–e1289. doi:10.1038/cddis.2014.253

56. Kim EJ, Sung JY, Lee HJ, Rhim H, Hasegawa M, Iwatsubo T, et al. Dyrk1A phosphorylates alpha-synuclein and enhances intracellular inclusion formation. J Biol Chem. 2006;281: 33250–33257.

57. Im E, Chung KC. Dyrk1A phosphorylates parkin at Ser-131 and negatively regulates its ubiquitin E3 ligase activity. J Neurochem. 2015;134: 756–768.

58. Mencacci NE, Isaias IU, Reich MM, Ganos C, Plagnol V, Polke JM, et al. Parkinson’s disease in GTP cyclohydrolase 1 mutation carriers. Brain. 2014;137: 2480–2492.

