## Supplementary Table 1 for "The Parkinson’s Disease GWAS Locus Browser"

| Locus Number | Risk Variant | LD Variant | Associated Disease | PMID | Frequency | P-value | r^2^ | D` |
| --- | --- | --- | --- | --- | --- | --- | --- | --- |
| 2 | rs6658353 | rs1801274 | Inflammatory bowel disease | 23128233 | 0.5018 | 2.00E-38 | 0.979216 | 0.991281 |
| 2 | rs6658353 | rs1801274 | Inflammatory bowel disease | 26192919 | 0.5018 | 9.00E-36 | 0.979216 | 0.991281 |
| 2 | rs6658353 | rs1801274 | Inflammatory bowel disease | 28067908 | 0.5018 | 9.00E-14 | 0.979216 | 0.991281 |
| 2 | rs6658353 | rs1801274 | Ulcerative colitis | 19915573 | 0.5018 | 2.00E-12 | 0.979216 | 0.991281 |
| 2 | rs6658353 | rs1801274 | Ulcerative colitis | 21297633 | 0.5018 | 2.00E-20 | 0.979216 | 0.991281 |
| 2 | rs6658353 | rs1801274 | Ulcerative colitis | 26192919 | 0.5018 | 1.00E-41 | 0.979216 | 0.991281 |
| 2 | rs6658353 | rs1801274 | Ulcerative colitis | 28067908 | 0.5018 | 2.00E-18 | 0.979216 | 0.991281 |
| 4 | rs823118 | rs823114 | Body mass index | 28892062 | 0.5572 | 4.00E-08 | 0.996607 | 0.999016 |
| 11 | rs1474055 | rs2390669 | Body mass index | 28892062 | 0.1282 | 6.00E-10 | 0.924287 | 0.992701 |
| 11 | rs1474055 | rs2390669 | Body mass index | 28892062 | 0.1282 | 2.00E-10 | 0.924287 | 0.992701 |
| 27 | rs26431 | rs28061 | Insomnia symptoms (never/rarely vs. usually) | 30804566 | 0.3011 | 1.00E-08 | 0.943624 | 0.980019 |
| 29 | rs4140646 | rs9380010 | Neuroticism | 30643256 | 0.2287 | 2.00E-15 | 0.877575 | 0.986445 |
| 61 | rs2904880 | rs4072402 | Body mass index | 31453325 | 0.6535 | 3.00E-28 | 0.913748 | 0.992749 |
| 61 | rs2904880 | rs4072402 | Body mass index variance | 31453325 | 0.6535 | 6.00E-12 | 0.913748 | 0.992749 |
| 62 | rs11150601 | rs12445568 | Neuroticism | 29255261 | 0.3814 | 4.00E-11 | 0.840179 | 0.920725 |
| 62 | rs11150601 | rs2199036 | Neuroticism | 30595370 | 0.3823 | 2.00E-09 | 0.840553 | 0.922019 |
| 64 | rs3104783 | rs3104778 | Insomnia symptoms (never/rarely vs. usually) | 30804566 | 0.4154 | 1.00E-08 | 0.992487 | 0.999599 |
| 69 | rs117615688 | rs113100008 | Neuroticism | 29942085 | 0.0619 | 2.00E-10 | 0.90233 | 0.997294 |
| 73 | rs1941685 | rs12605642 | Insomnia | 30804565 | 0.4886 | 2.00E-09 | 0.961045 | 0.997581 |
| 73 | rs1941685 | rs10460051 | Neuroticism | 29942085 | 0.5026 | 3.00E-09 | 0.80383 | 0.909439 |
| 73 | rs1941685 | rs11876492 | Neuroticism | 30595370 | 0.4864 | 2.00E-10 | 0.827711 | 0.9407 |
| 73 | rs1941685 | rs1493914 | Neuroticism | 30643256 | 0.5138 | 2.00E-11 | 0.804814 | 0.927574 |

**Supplementary Table 1. Select known GWAS variants in linkage disequilibrium with PD risk variants.** List of variants in LD with PD risk variants with r^2^ > 0.8, D` > 0.9 and P-value < 5E-08. Frequency included is for non-Finnish European populations. Only variants with phenotypes for body mass index, neuroticism, insomnia, ulcerative colitis, and inflammatory bowel disease are included.
